# ADARs mediate distinct RNA editing activity and gene regulation in the *Caenorhabditis elegans* germline

**DOI:** 10.1101/2025.10.16.682908

**Authors:** Emily A. Erdmann, Heather A. Hundley

## Abstract

Tissues rely on unique landscapes of gene regulation to allow the organism to correctly develop, function, and respond to changes. One component of these gene regulatory networks is RNA Binding Proteins (RBPs) which bind and modify RNA molecules leading to changes in the cellular fate of transcripts. The Adenosine DeAminase acting on RNA (ADAR) family of RBPs modify RNAs by catalyzing the deamination of adenosine (A) to inosine (I), known as A-to-I RNA editing. Prompted by recent evidence that ADARs play important roles in germline biology, we profiled editing activity of the A-to-I editing enzyme ADR-2 on transcripts in the *Caenorhabditis elegans* germline. These analyses revealed that many germline editing events are distinct from editing events in other tissues; however, the previously described role of the inactive deaminase ADR-1 in regulating editing activity by ADR-2 is conserved in the germline. We find that complete loss or misregulation of editing has little effect on the expression of edited transcripts within the germline; however, loss of ADARs results in the misexpression of several unedited germline transcripts. Intriguingly, further investigation reveals that these expression changes are buffered at the translational level. In all, the results of this study suggest that ADARs show unique activity in the *C. elegans* germline and that compensatory mechanisms exist to lessen the immediate consequences of loss of ADAR function within the germline.

## Introduction

RNA-level regulation of gene expression is essential in coordinating proper development and organismal function. In particular, RNA regulation is important in tissue diversification (Franks et al. 2017). One mechanism of RNA regulation important for proper tissue development is RNA modifications, by which RNAs are chemically altered to affect their cellular fate (Porat 2025).

Adenosine-to-Inosine (A-to-I) RNA editing is one of the most abundant RNA modifications in metazoans (Zhang et al. 2023) and is catalyzed by the Adenosine DeAminase acting on RNA (ADAR) family of RNA binding proteins (Datta et al. 2023). A-to-I editing can have various effects on transcript fate (Eisenberg and Levanon 2018). As inosine is read as guanosine by the translational machinery, editing in coding regions of transcripts can lead to changes in codons, resulting in changes in the amino acid sequence of the encoded peptide (Lewis 2023). While this “recoding” editing can have important impacts, A-to-I editing is abundant within noncoding regions of transcripts (Bazak et al. 2014). Noncoding editing primarily affects RNA processing and regulation and can lead to changes in gene expression which ultimately result in physiological changes at the cell, tissue, and organismal levels (Rueter et al. 1999; Yang et al. 2006; Lev-Maor et al. 2007; Scadden 2007; Yang et al. 2017; Hsiao et al. 2018; Liu et al. 2024). A-to-I editing patterns and frequency are known to differ throughout developmental stages and tissues (Lev-Maor et al. 2007; Ekdahl et al. 2012; Tan et al. 2017; Rajendren et al. 2021). As such, tissue and cell type-specific studies have proven useful in identifying novel molecular and biological consequences of editing (Fei et al. 2016; Deffit et al. 2017; Chen et al. 2024).

In addition to A-to-I editing, ADARs are double-stranded RNA (dsRNA) binding proteins and can target dsRNAs ranging from small RNAs and their precursors to longer transcripts containing intramolecular duplexes (Marônek et al. 2025). dsRNA binding by ADARs can impact transcripts by competing with other RNA binding factors for targets or by recruiting other RNA binding factors to targets (Karlström et al. 2024; Mahapatra et al. 2025). These interactions can ultimately lead to changes in transcript processing and stability. As such, several editing-independent effects of ADARs have also been shown to play important roles in tissue development and functions (Capshew et al. 2012; Cho et al. 2018; Deng et al. 2020; Erdmann et al. 2024).

Due to the severe neural defects originally reported upon loss of ADARs in mammals (Burns et al. 1997; Higuchi et al. 2000), tissue-specific studies have mainly focused on the nervous system (Liscovitch et al. 2014; Deffit et al. 2017; Sapiro et al. 2019; Rajendren et al. 2021). However, transcriptomic studies have identified ADAR expression and editing events across a wide range of human tissues (Tan et al. 2017). Recently, emerging evidence from mammalian studies has pointed to biological roles for ADARs in germline development and function (Snyder et al. 2020; Nelson et al. 2022). While previous studies have identified editing events in germline tissues, the molecular consequences of ADARs on germline transcripts remain largely unknown.

The nematode *Caenorhabditis elegans* is an ideal model system for assessing the impacts of ADARs as null mutants are viable (Tonkin et al. 2002). While A-to-I editing and ADAR function has been studied extensively at the whole animal level in *C. elegans*, (Washburn et al. 2014; Whipple et al. 2015; Zhao et al. 2015; Rajendren et al. 2018; Ganem et al. 2019; Eliad et al. 2024; Schiksnis et al. 2024) and in neural tissues (Deffit et al. 2017; Rajendren et al. 2021), there is little understanding of ADAR activity in the germline. Given emerging evidence that ADR-2, the sole A-to-I editing enzyme in *C. elegans*, is expressed widely in germline tissues (Eliad et al. 2024) and plays roles in regulating germline processes (Reich et al. 2018; Fischer and Ruvkun 2020; Erdmann et al. 2024), this study aimed to investigate how ADR-2 regulates transcripts in the *C. elegans* germline. Using an unbiased, high-throughput approach, we profile the molecular targets of germline ADARs and characterize the consequences of ADAR function on the expression and translation of these transcripts.

## Results and Discussion

### ADARs are expressed throughout the C. elegans adult germline

In a previous study, *adr-2* mRNA expression was found to be significantly reduced in young adult neural cells compared to L1 neural cells; however, the total *adr-2* mRNA expression in the animal remained unchanged (Rajendren et al. 2021), begging the question of where most *adr-2* expression is taking place in young adult animals. As the onset of young adulthood corresponds to reproductive maturity in *C. elegans* (Pazdernik and Schedl 2013), we were curious whether this expression could be coming from the germline. To determine the contribution of the germline to total ADAR expression in young adult animals, mRNA expression of the active deaminase *adr-2* and the other *C. elegans* ADAR family member, the inactive deaminase *adr-1* (Washburn et al. 2014), was measured in wild-type animals and *glp-4(bn2)* temperature sensitive mutant animals which fail to develop germ cells when raised at a restrictive temperature (25°C) (Beanan and Strome 1992). Genomic deletion mutants of *adr-1* and *adr-2* were included as negative controls. Strikingly, germline-lacking *glp-4(bn2)* animals show a significant decrease in *adr-2* expression compared to germline-containing wild-type animals (**Figure 1A**, left), suggesting that the majority of *adr-2* mRNA expression in young adult animals is within the germline. Similar results were seen for *adr-1*, with *glp-4(bn2)* germline-lacking animals showing similar *adr-1* mRNA levels to those detected in *adr-1(-)* animals, a significant decrease compared to wild-type, germline-containing animals (**Figure 1A**, right). Together, these data suggest that ADARs are abundantly expressed at the mRNA level in the germline of young adult animals. However, as mRNA expression does not always scale with protein expression, and as many germline transcripts are translationally repressed and stored for use in the early embryo (Quarato et al. 2021), it is important to confirm whether ADARs are also expressed at the protein level in the germline.

**Figure 1:**
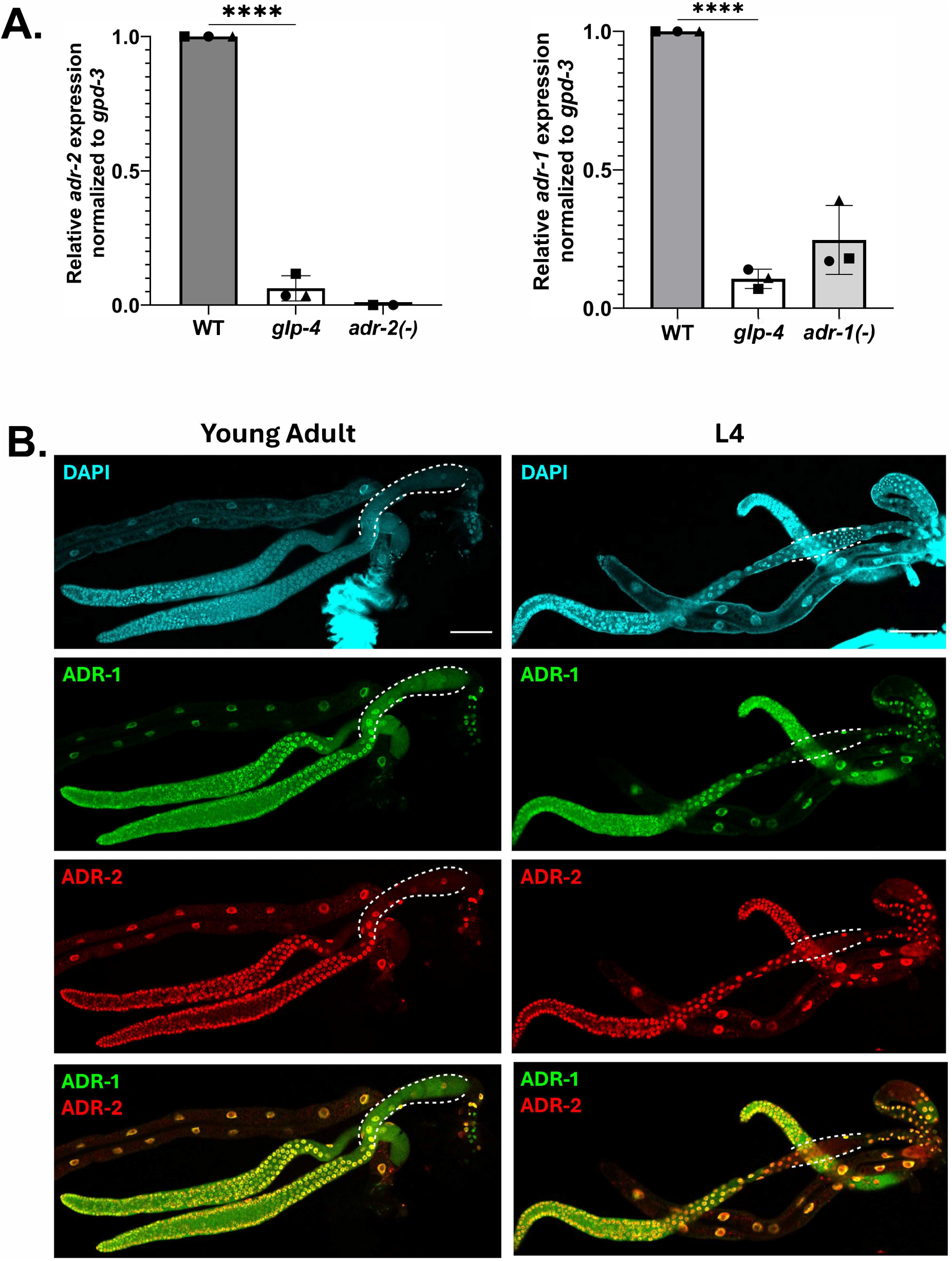
ADARs are abundantly expressed in the young adult *C. elegans* germline. A) Relative mRNA expression of *adr-2* (left) and *adr-1* (right) in wild-type, germline-lacking *glp-4* temperature-sensitive mutant, and *adr* null mutant young adult animals raised at the restrictive temperature of 25°C. *adr* expression is normalized to the housekeeping gene *gpd-3*. Data represents three biological replicates. Significance determined via one-way Anova, **** indicates p ≤ 0.0001. B) Representative confocal images of young adult (left) and L4 (right) extruded germlines stained for DAPI (top, cyan), V5::ADR-1 (top middle, green), and 3xFLAG::ADR-2 (bottom middle, red). Bottom panel shows signals of ADR-1 and ADR-2 panels merged. Dotted lines denote proximal germline region containing oocytes (young adult) or sperm (L4). Each image includes two gonad arms and the intestine from the same animal. Scale bars = 50 μm.

Recently, we have shown that ADR-2 protein is expressed in germ nuclei throughout gametogenesis, and in mature oocytes but not in mature sperm (Erdmann et al. 2024). To determine where ADR-1 is expressed in the germline and whether it correlates with ADR-2 expression, germlines were extruded from young adult and L4 animals and immunostaining was performed for both V5-tagged ADR-1 and 3xFLAG-tagged ADR-2. ADR-1 shows a similar expression pattern to that of ADR-2, with expression in germ nuclei throughout gametogenesis and in oocytes (**Figure 1B**, left) but not in sperm (**Figure 1B**, right). While both ADARs show a strong nuclear signal, ADR-1 also appears to be distributed throughout the cytoplasm (**Figure 1B**) while, consistent with previous reports, ADR-2 appears more restricted to the nucleus (Eliad et al. 2024; Erdmann et al. 2024).

### Profiling A-to-I RNA editing activity on germline transcripts

Given our results that both ADR-2 and ADR-1 are expressed throughout the germline, we sought to determine whether ADARs are active in catalyzing and regulating A-to-I RNA editing in the germline. To profile A-to-I editing in the young adult *C. elegans* germline, germline RNA sequencing was performed. Sequencing libraries were prepared from dissected germline samples from wild-type and *adr-2(-)* animals and sequencing reads were aligned to a reference genome. To identify editing sites in germline transcripts, we used SAILOR, a bioinformatic program which identifies high-confidence A-to-I editing sites (Deffit et al. 2017; Kofman et al. 2023).

Putative A-to-I editing sites were restricted to sites with a reproducible confidence level ≥ 75% from SAILOR in all three biological replicates (1098 putative editing sites) of the wild-type germline samples. As *adr-2(-)* animals lack A-to-I editing (Tonkin et al. 2002; Washburn et al. 2014), the *adr-2(-)* RNA-seq data serves as a control for the computational analysis. Consistent with this, SAILOR analysis on the RNA-seq from *adr-2(-)* germlines identified 86 sites that were also identified in the wild-type animals, which were in turn removed as false positives. These analyses identified a final list 1012 high-confidence A-to-I RNA editing sites in 71 germline transcripts (**Table S1**). Similar to editing site analysis from adult whole worms (Schiksnis et al. 2024) and adult neural cells (Rajendren et al. 2021), the majority of germline editing sites were located in non-coding regions of transcripts, primarily the 3’ untranslated region (UTR) (**Figure 2A, Table S1**).

**Figure 2:**
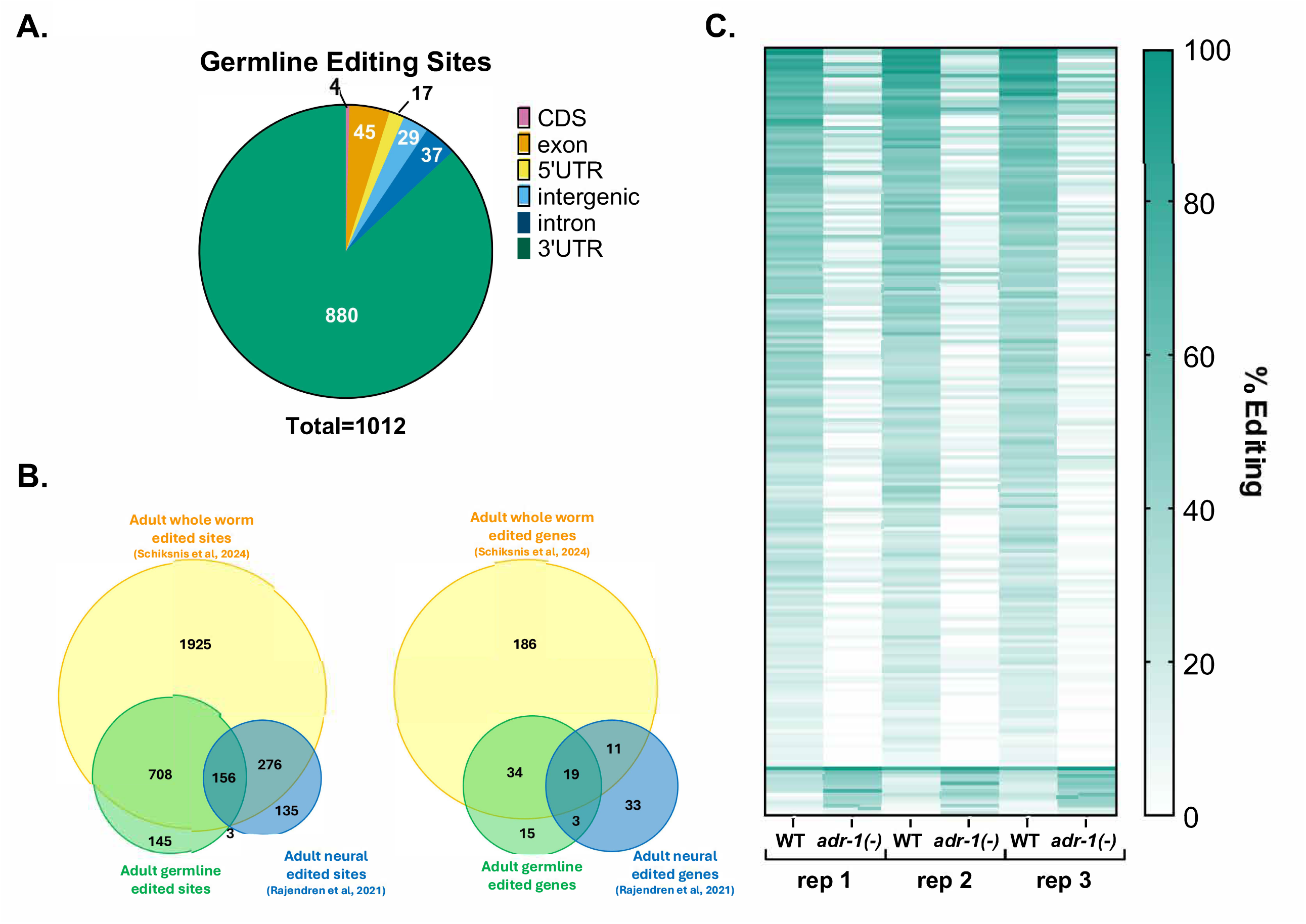
ADR-2 edits distinct targets in the germline and is regulated by ADR-1. A) Genic distribution of high-confidence germline editing sites identified in RNA-seq data from germlines of wild-type animals. B) Comparison of edited sites (left) and edited genes (right) between adult germline samples (this study, green), adult neural samples (Rajendren et al. 2021)(blue), and adult whole worm samples (Schiksnis et al. 2024)(yellow). C) ADR-1-regulated germline editing sites. Percent editing at individual germline editing sites for three independent biological replicates of RNA-seq data from wild-type and *adr-1(-)* germlines.

Comparing our list of germline editing sites to a previously published dataset from adult worms (Schiksnis et al. 2024), we found a strong overlap in edited sites and edited genes, as would be expected because the adult samples were from the entire animal, and thus, include the germline (**Figure 2B, Table S1**). To get a better idea of how germline editing compares to other tissues, we also compared our list of germline editing sites to a previously published dataset from adult *C. elegans* neural cells (Rajendren et al. 2021). While there was some overlap between the neural and germline editomes (159 sites in 22 genes), the majority of neural and germline editing sites were mutually exclusive, with 411 sites and 44 genes exclusively identified in the neural editome and 853 sites and 49 genes exclusively identified in the germline editome (**Figure 2B, Table S1**). One possibility for the identification of unique germline edited genes is that the genes are exclusively expressed in the germline and thus unable to be edited in other tissues. To determine whether tissue-specific expression was driving identification of germline-specific editing, we assessed the expression patterns of the 49 germline-specific edited genes and found that all of these genes are also expressed in neurons in young adult animals (Ghaddar et al. 2023) (**Table S1**). These data demonstrate that while some A-to-I editing sites and edited genes are conserved across tissues, there are also tissue-specific differences in editing patterns, suggesting differential regulation of ADAR activity across tissues and potentially unique biological consequences of editing in different tissues.

ADR-1 has been shown to regulate A-to-I editing by ADR-2 in neural tissues (Washburn and Hundley 2016; Rajendren et al. 2021). Due to the differences in ADR-2 editing patterns seen between neural and germline tissues, we wanted to determine whether ADR-1 regulates editing of germline transcripts. We assessed editing levels for germline edited sites in wild-type and *adr-1(-)* germlines by performing variant calling on germline RNA-seq data. Variants mapping to our 1012 high-confidence editing sites were selected for further analysis. To ensure identification of only robust changes in editing, regulated sites were defined as those with read coverage of greater than 10 reads in both wild-type and *adr-1(-)* sequencing data and exhibiting a greater than |5%| difference in editing between wild-type and *adr-1(-)* for all three biological replicates. Upon applying these parameters, 205 germline editing sites were designated as ADR-1-regulated (**Figure 2C, Table S2**). For the vast majority of these sites (192/205), editing decreased upon loss of *adr-1*, suggesting that ADR-1 promotes ADR-2 activity at these sites in wild-type germlines. The remaining 13 sites showed an increase in percent variant reads in *adr-1(-)* germlines compared to wild-type, suggesting ADR-1 inhibits editing at these sites in wild-type germlines. Taken together, these data suggest that, as previously shown in other tissues (Washburn et al. 2014; Rajendren et al. 2021), ADR-1 regulates A-to-I editing by ADR-2 in the germline, primarily facilitating editing but also inhibiting editing at some sites.

### ADR-2 regulates expression of ribosomal RNA

To investigate whether A-to-I editing can affect mRNA expression of germline transcripts, we performed differential gene expression analysis between wild-type and *adr-2(-)* germlines. Edited genes show minimal changes in expression between wild-type and *adr-2(-)* (**Figure 3A, Table S3**), with only one edited gene (*gid-7*) showing a significant change in expression (padj ≤ 0.05, log2 fold change ≥ 0.5). This suggests that A-to-I editing does not directly impact germline mRNA expression of most edited transcripts in *C. elegans*.

**Figure 3:**
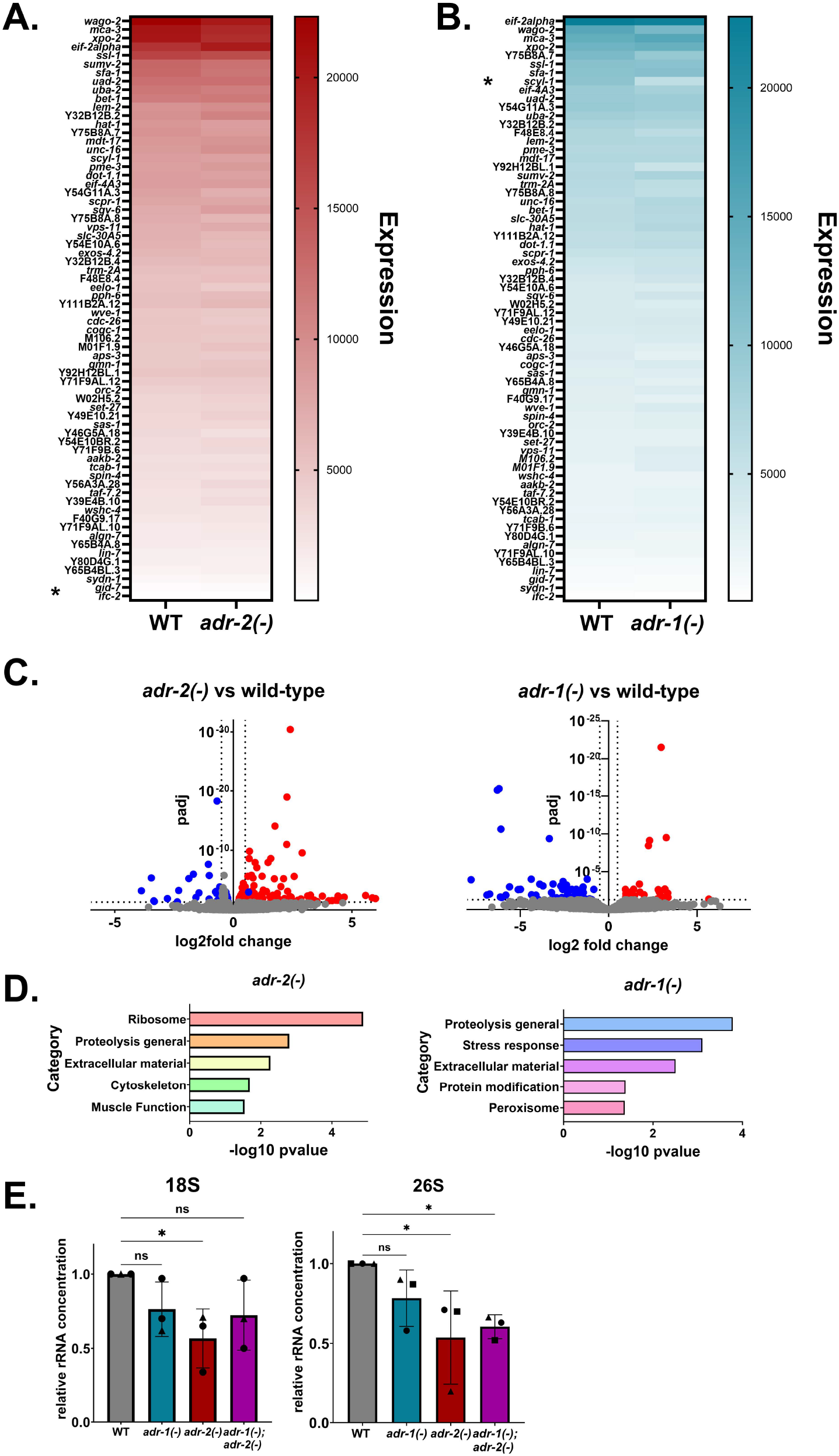
ADARs regulate germline gene expression. A-B) Expression of edited genes in *adr-2(-)* germlines compared to control (WT) germlines (A) and *adr-1(-)* germlines compared to control (WT) germlines (B). Data represents expression values from DESeq analysis averaged across three biological replicates. * indicates significant change in expression (padj ≤ 0.05, log2 fold change ≥ 0.05). C) Differentially expressed genes in *adr-2(-)* germlines compared to wild-type (left) and *adr-1(-)* germlines compared to wild-type (right). Gray points denote genes with no significant change in expression, red points denote significantly upregulated genes (padj ≤ 0.05, log2 fold change ≥ 0.5), blue points denote significantly downregulated genes (padj ≤ 0.05, log2 fold change ≤ 0.5). Two outlying genes were left out of *adr-2(-)* figure for scale: *adr-2* (log2 fold change= -6.13, padj= 2.23×10^-241^) and *cls-3* (log2 fold change= 3.77, padj= 1.40×10^-123^). D) Gene ontology analysis of genes misregulated in *adr-2(-)* germlines compared to wild-type (left) and *adr-1(-)* germlines compared to wild-type (right). E) Tapestation measurements of 18S and 26S rRNA in wild-type, *adr-1(-), adr-2(-)*, and *adr-1(-);adr-2(-)* germlines. rRNA concentrations are normalized to a spike-in RNA of known concentration. Data represents three biological replicates. Significance determined via one-way Anova, ns indicates p > 0.05, * indicates p ≤ 0.05.

While loss of editing did not show significant impacts on germline gene expression, previous studies have shown that misregulated editing can have more severe impacts than loss of editing altogether (Ganem et al. 2019). As such, we were interested in whether loss of regulation by ADR-1 affects the expression of edited transcripts. To investigate this, differential gene expression analysis was performed for germlines isolated from *adr-1(-)* animals compared to those isolated from wild-type animals. Similar to what was observed upon loss of editing, loss of *adr-1* resulted in minimal changes in gene expression across edited transcripts, with only one edited transcript (*scyl-1*) showing significant changes in expression (padj ≤ 0.05, log2 fold change ≥ 0.5). Of note, ADR-1-regulated editing sites (**Figure 3B, Table S3**) were not found in *scyl-1* in our analysis, suggesting that any changes in its expression upon loss of *adr-1* do not result from *adr-1*-dependent changes in editing.

While aberrant editing activity does not appear to affect the expression of germline-edited genes, loss of either *adr-2* or *adr-1* resulted in misregulation of several unedited genes. In the absence of *adr-2*, 93 genes were significantly upregulated compared to wild-type and 34 genes were significantly downregulated compared to wild-type (**Figure 3C, left, Table S3**). Loss of *adr-1* had a slightly lesser impact on gene expression than *adr-2*, with 25 genes significantly upregulated compared to wild-type and 60 genes significantly downregulated compared to wild-type (**Figure 3C, right, Table S3**). Notably, there was little overlap between the genes misregulated upon loss of *adr-2* and *adr-1*, with only 5 genes misregulated in both datasets. To identify any common themes in the genes regulated by *adr-2* and *adr-1* in the germline, gene ontology analysis was performed using a *C. elegans* specific program, WormCat (Holdorf et al. 2020) (**Figure 3D, Table S3**). This analysis showed that loss of *adr-2* and loss of *adr-1* both affected expression of genes involved in proteolysis and extracellular material; both of these categories have been shown to be affected by loss of ADR-2 in the nervous system as well (Mahapatra et al. 2023), and extracellular material was shown to be affected by loss of ADARs in whole worms (Dhakal et al. 2024). Uniquely, the genes misregulated by loss of *adr-2* was most highly enriched for genes associated with ribosomes (8/127 genes; pvalue = 1.31e-5), including ribosomal subunit proteins as well as ribosomal RNAs (rRNAs) (**Figure 3D, left**).

Because the preparation of the samples for sequencing included poly-A selection, which should select against rRNAs, we sought to independently test whether the rRNA reduction observed in the *adr-2(-)* germline RNA-seq was due to loss of *adr-2* or simply an artifact of differential poly-A selection efficiencies across samples. Additionally, we measured rRNA levels in *adr-1(-)* and *adr-1(-);adr-2(-)* animals to determine whether this was a general effect of loss of ADARs. A separate set of germline RNA samples was collected from wild-type, *adr-1(-), adr-2(-)*, and *adr-1(-);adr-2(-)* animals. The total RNA for each sample was analyzed using Agilent Tapestation electrophoresis. For all samples, a spike-in RNA of known size and concentration was added to normalize for differences in RNA isolation efficiency across samples. Agilent software was used to determine the concentrations of 18S and 26S rRNA in each sample relative to the concentration of spike-in RNA. While rRNA levels were unaffected in *adr-1(-)* animals, both 18S and 26S rRNA populations were significantly reduced in *adr-2(-)* germline samples compared to wild-type germline samples (**Figure 3E**). Only 26S rRNA was significantly reduced in *adr-1(-);adr-2(-)* germline samples. These data suggest that the reduction in expression of rRNAs seen in our sequencing analysis represents an effect on germline rRNA populations upon loss of *adr-2*.

### Altered gene expression upon loss of adr-2 are buffered at the translational level

The reduction of rRNA levels observed in *adr-2(-)* germlines raised the question of whether translation of germline transcripts were altered in animals lacking *adr-2*. To profile translational activity in a germline-specific manner, relative ribosome association of germline transcripts between wild-type and *adr-2(-)* animals was examined using a tissue-specific Translating Ribosome Affinity Purification (TRAP) method (Nousch 2020) followed by high-throughput sequencing. Briefly, strains expressing an integrated, epitope tagged ribosome protein (RPL-4::FLAG) under a germline specific promoter (*mex5p*) were synchronized and grown to young adulthood. Animals were treated with cycloheximide to induce ribosome stalling and lysates were generated. Immunoprecipitation was performed using FLAG antibody-conjugated magnetic beads to pull down ribosomes and their associated RNAs.

To validate the tissue-specificity of this method, ribosome-bound RNAs were isolated and enrichment of representative germline-expressed and soma-expressed genes in the immunoprecipitated samples was quantified using reverse transcription followed by qRT-PCR. Significant enrichment over a negative control (wildtype/non-tagged strain) was observed in the germline RPL-4::FLAG strain for the germline-expressed gene *glh-1*, suggesting that germline-translated transcripts are pulled down with this method. Conversely, no significant enrichment over the negative control was seen for the soma-expressed gene *elt-2* (**Figure S1**). These data demonstrate that the TRAP method is specific for germline ribosomes and their associated RNAs.

To identify the effects of *adr-2* on translation, the germline *RPL-4::FLAG* construct was crossed into an *adr-2* deletion mutant (*adr-2(ok735)*). Importantly, *rpl-4* is not among the ribosome components downregulated by loss of *adr-2*, and *adr-2(-)* animals showed similar expression of RPL-4::FLAG compared to wild-type (**Figure S2**). Germline TRAP was performed on three biological replicates each of wild-type and *adr-2(-)* animals, and cDNA libraries were prepared and sequenced from ribosome-bound RNA fractions (**Figure S2**). Next, ribosome association of germline transcripts was compared between wild-type and *adr-2(-)* animals. This analysis found minimal change in ribosome association between samples for most transcripts, with only 3 genes showing significant changes (padj ≤ 0.05) in ribosome association between wild-type and *adr-2(-)*, including 1 gene with decreased association and 2 genes with increased association (**Figure 4A**). However, these results are intriguing when compared to our mRNA expression experiment described earlier (**Figure 3C, Table S3**), which showed that over 100 genes are misregulated at the mRNA expression level in *adr-2(-)* germlines compared to wild-type. The lack of a similar change in ribosome association for these misregulated genes suggests that the changes in mRNA level caused by loss of *adr-2* do not lead to differences in association of the mRNAs with the ribosome (**Figure 4A-B**). As such, it is unlikely that these changes in mRNA expression are mirrored at the protein expression level. Additionally, this suggests, as seen in other germline studies, the presence of secondary regulation which can buffer changes in mRNA expression to maintain proper translational output (Rochester et al. 2022). This finding may also explain why loss of *adr-2* alone, despite causing significant changes in gene expression at the mRNA level (**Figure 3C, Table S3**), does not cause any obvious defects in germline development or function. While studies of germline gene expression regulators have demonstrated the presence of redundant and overlapping regulatory pathways that act as fail-safes to ensure proper germline function (Vanden Broek et al. 2022), future studies are needed to understand the mechanism of regulation at play here.

**Figure 4:**
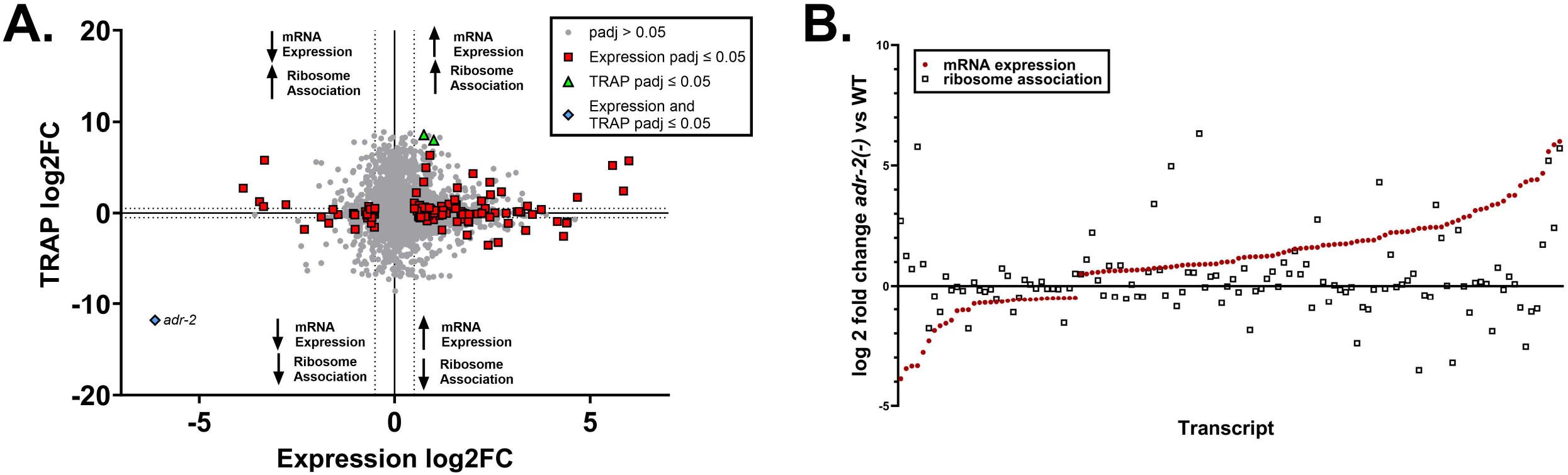
ADR-2-dependent gene expression changes are buffered at the translational level. A) Quadrant scatter plot showing the fold change (log2) of germline expression (x axis) and TRAP ribosome association (y axis) in *adr-2(-)* germlines compared to wild-type. Gray points denote genes with no significant change in either analysis. Red squares indicate a significant change in mRNA expression only (DESeq2 padj ≤ 0.05, log2 fold change |≥ 0.5|) in *adr-2(-)* compared to wild-type, green triangles indicate a significant change in ribosome association only (DESeq2 padj ≤ 0.05, log2 fold change |≥ 0.5|) in *adr-2(-)* compared to wild-type. Blue diamond indicates a significant change in both expression and ribosome association (*adr-2* only). DESeq2 analyses performed with three biological replicates. Dotted lines represent log2 fold change significance cutoff. B) Log2 fold change mRNA expression (red circles) and log2 fold change germline ribosome association (black squares) in *adr-2(-)* germlines compared to wild-type for genes showing significant changes in mRNA expression (DESeq2 padj ≤ 0.05, log2 fold change |≥ 0.5|).

In sum, we demonstrate in this study that A-to-I RNA editing occurs at over 1000 sites across 71 germline transcripts. Interestingly, we see minimal changes in mRNA expression and translational status of edited transcripts in the absence of *adr-2*. While it is possible that editing has no effect on these transcripts under normal conditions, it is also possible that the effects of editing on these transcripts are not evident within the adult germline, where these experiments were performed. In addition to executing gametogenesis, a major function of the germline is to prepare for early embryonic development. As such, a cache of maternal transcripts are transcribed during gametogenesis and stored for use in early embryonic development before the onset of zygotic transcription (Rajyaguru and Parker 2009). Thus, editing of maternal transcripts in the germline may affect their fate in the embryo. While many of the germline edited transcripts identified in this study are known maternal transcripts (Quarato et al. 2021) (**Table S1**), further studies are needed to determine whether editing events are present on inherited early embryonic transcripts. It is also possible that the whole-germline nature of our studies has obscured effects on edited genes that may be restricted to specific cell types or stages in the germline. Single-cell studies may shed light on these cell-type specific interactions and their potential effects on germline development and function.

Additionally, we demonstrate that *adr-2* affects the expression of several ribosome components and rRNAs, leading to an overall reduction in both 18S and 26S rRNA concentrations. Studies of ribosome composition have revealed that ribosomes are highly heterogeneous, consisting of different sets of ribosomal proteins and rRNAs (Gay et al. 2022). Further studies have revealed that this heterogeneity can have functional consequences, with ribosomes of different composition preferentially translating different subsets of transcripts (Shi et al. 2017; Norris et al. 2021). Ribosome compositions have been shown to vary between tissues, developmental timepoints, and environmental conditions to facilitate unique translational regulation with functional consequences (Hopes et al. 2022; Li et al. 2022; Milenkovic and Novoa 2025). As such, the regulation of ribosome components by ADR-2 in the germline may contribute to germline-specific patterns of ribosome composition that ultimately regulate the germline translational landscape. Future studies are needed to elucidate the molecular relationship between ADR-2, ribosome component expression, and germline translational regulation, as well as the downstream consequences of ADR-2-dependent gene regulation on germline development and function.

## Materials and Methods

### Worm strains and maintenance

All worm strains were maintained at 20°C on nematode growth media (NGM) seeded with *Escherichia coli* OP50. Worms were thawed regularly from frozen stocks to minimize effects of accumulated random mutations. The following previously generated strains were used in this study: Bristol Strain N2, SS104 (*glp-4(bn2)*) (Beanan and Strome 1992), BB19 (*adr-1(tm668)*) (Hundley et al. 2008), BB20 (*adr-2(ok735)*) (Hundley et al. 2008), BB21 (*adr-1(tm668);adr-2(ok735)*) (Hundley et al. 2008), EV484 (efls155[*mex-5p*::*rpl-4*::FLAG::*tbb-2* 3’UTR + *Cbr-unc-199*(+)]II) (Nousch 2020), HAH36 (V5::*adr-1*;3xFLAG::*adr-2*) (Dhakal et al. 2024), and HAH61 (3xFLAG::*adr-2*) (Dhakal et al. 2024). Strains generated in this study include: HAH87 (efls155[*mex-5p*::*rpl-4*::FLAG::*tbb-2* 3’UTR + *Cbr-unc-199*(+)]II), HAH88 (*adr-2(ok735)*; efls155[*mex-5p*::*rpl-4*::FLAG::*tbb-2* 3’UTR + *Cbr-unc-199*(+)]II).

Crossed strains were made by placing 10-15 males and 1 hermaphrodite on mating plates (NGM plates seeded with a small spot of *E. coli* OP50 in the center) and genotyping was performed for the F1 progeny and F2 progeny using primers mentioned in **Table S5**. The specific crosses performed included: creation of HAH87 and HAH88 by crossing EV484 hermaphrodites to BB20 males.

### Bleaching

Synchronized animals were obtained by bleaching gravid adult animals with a solution of 5M NaOH (33%) and Sodium Hypochlorite (Fisher Scientific) (66%). After the solution was added, animals were incubated on a shaker at 20°C for 7 minutes and then spun down to collect embryos. Embryos were washed with 1x M9 buffer (22 mM KH_2_PO_4_, 42.3 mM Na_2_HPO_4_, 85.6 mM NaCl, 1 mM MgSO_4_) solution thrice and incubated overnight in 1x M9 buffer at 20°C. The next day, hatched L1 worms were spun down and washed again with 1x M9 buffer thrice.

### Quantitative Real-Time PCR

RNA was extracted using standard Trizol-Chloroform extraction followed by treatment with TURBO DNase (Ambion) and cleanup with the RNeasy Extraction Kit (Qiagen). Reverse transcription was performed on total RNA using random hexamer primers, oligo dT, and Superscript III (Thermo Fisher). Gene expression was determined using KAPA SYBR FAST Master Mix (Roche) and gene-specific primers (**Table S6**) on a Thermo Fisher Quantstudio 3 instrument. The primers designed for qPCR spanned an exon-exon junction to prevent detection of genomic DNA in the samples. For each gene analyzed, a standard curve of 8 to 10 samples of 10-fold serial dilutions of the amplified product were used to generate a standard curve of cycle threshold versus the relative concentration of amplified product. Standard curves were plotted on a logarithmic scale in relation to concentration and fit with a linear line. Fit (r^2^) values were around 0.99 and at least 7 data points fell within the standard curve. Each cDNA measurement was performed in 3 technical replicates, and each experiment was performed in 3 biological replicates.

### Germline Immunostaining & Imaging

Germlines were extruded from synchronized L4 or day-one adult HH36 animals. Germlines were fixed with 1% paraformaldehyde and methanol and permeabilized with 0.2% Triton X-100 (Sigma-Aldrich). Germlines were stained with anti-V5 (Cell Signaling Technology) and anti-FLAG (Sigma-Aldrich) primary antibodies, followed by Alexa Fluor 594 (Thermo Fisher Scientific), Alexa Fluor 488 (Thermo Fisher Scientific), and DAPI (4′,6-diamidino-2-phenylindole) (Thermo Fisher Scientific). Germlines were mounted in Vectashield (VectorLabs) and imaged using a Leica SP8 Scanning Confocal microscope (Indiana University Light Microscopy Imaging Center). Images were processed using ImageJ.

### Germline Sample Collection for RNA-sequencing

Adapted from (Campbell and Updike 2015). Synchronized young adult (60 hour post egg-lay) animals were used for germline dissection. Animals were picked into egg buffer (118 mM NaCl, 48 mM KCl, 2 mM CaCl_2_, 2 mM MgCl_2,_ 25 mM HEPES, pH 7.3 adjusted osmolarity to 340 mOsm with sucrose) containing levamisole (2.5mM). Animals were cut at the pharynx to allow for extrusion of the gonad. Germlines were collected via mouth aspiration using a pulled glass capillary coated with Sigmacote (Sigma-Aldrich). For germline RNA samples, germlines were dispensed into 400 ul Trizol on ice then frozen in liquid nitrogen. RNA was extracted using standard Trizol-Chloroform extraction followed by treatment with TURBO DNase (Ambion) and cleanup with the RNeasy Extraction Kit (Qiagen).

### RNA sequencing library preparation

For both the germline RNA-seq experiments and the TRAP-seq experiment: DNase-treated total RNA was subjected to two rounds of Poly-A selection using magnetic oligo(dT) beads (Invitrogen). Libraries were created from entire poly A-selected RNA samples using the KAPA Stranded RNA-seq Library Prep kit (Roche); cleanup steps were performed using AMPure XP beads (Beckman Coulter). Library fragment size distribution (200-900 bp) was determined via Tapestation electrophoresis (Agilent). Libraries for each of three biological replicates per condition were pooled by concentration and sequenced on a NextSeq 2000 (75 cycle) flow cell at the Indiana University Center for Genomics and Bioinformatics. The quality of the sequencing reads obtained was checked using FASTQC (version 0.12.1), the results of which are summarized in **Table S6**.

### Bioinformatic Analysis of Germline RNA-sequencing Data

Reads were aligned to a reference genome (*C. elegans* PRJNA13758 WS275) using STAR (version 2.7.10b). Percent uniquely mapped reads for each sample are listed in **Table S5**. A-to-I editing site identification was performed using SAILOR (Deffit et al. 2017; Kofman et al. 2023). Each biological replicate was analyzed separately. Editing sites called at ≥ 75% confidence in all three replicates of wild-type germlines were selected as high-confidence editing sites. Editing sites also called in *adr-2(-)* germlines (regardless of confidence) were removed as false positives (**Figure S3**). Variant analysis was performed using BCFtools mpileup (Danecek et al. 2021). Differential gene expression analyses were performed in R with DESeq2 (Love et al. 2014). Gene ontology enrichment analyses were performed with WormCat (Holdorf et al. 2020). Full code for these analyses is available at: https://github.com/emierdma/GSF3037.

### Germline sample collection and rRNA measurement

Germlines were dissected for RNA isolation as described above. An equal amount of a spike-in RNA of known concentration (*C. elegans lam-2* synthesized via *in vitro* transcription, adapted from (Rajendren et al. 2018)) was added to each germline sample, then RNA was extracted using standard Trizol-Chloroform extraction followed by treatment with TURBO DNase (Ambion) and cleanup with the RNeasy Extraction Kit (Qiagen). Total RNA was run on the Agilent Tapestation using High Sensitivity RNA ScreenTape (Agilent). Peaks for the spike-in RNA (160 nt), 18S rRNA (∼1000 nt), and 26S rRNA (∼2000 nt) were automatically assigned by the Tapestation software, with adjustments as needed. Concentrations for spike-in, 18S rRNA, and 26S rRNA were calculated by the Tapestation software.

### Translating Ribosome Affinity Purification (TRAP)

Animals were synchronized via bleaching and allowed to grow to young adulthood. Young adult animals were washed from plates with 1x M9 buffer and incubated for 20 minutes without food to allow for digestion of OP50. Animals were then washed once in extract buffer (50 mM HEPES, 70 mM K-Acetate, 5 mM Mg-Acetate, 10% Glycerol, 1% NP-40, pH 7.4) with added EDTA-free protease inhibitor (Roche) and cycloheximide (100 μg/ml) and resuspended in an equal volume of concentrated extract buffer with protease inhibitor and cycloheximide before being dripped into liquid nitrogen to make pellets. Pellets were ground to a fine powder and allowed to thaw, then centrifuged to generate a clear lysate. Input lysate was taken and either boiled with 6x SDS buffer for 5 minutes for western blotting or frozen in 400 ul Trizol for RNA isolation. For each immunoprecipitation (IP), 200 ug of lysate was added to 25 ul of prewashed FLAG magnetic resin along with 200 ul of concentrated extract buffer with cycloheximide and brought up to 1 ml volume with extract buffer with no added detergents. Lysates were incubated with resin for 2 hours at 4°C, then washed thrice with KCl wash buffer (10 mM HEPES pH 7.4, 150 mM KCl, 5 mM MgCl_2_, 1% NP-40) with added cycloheximide (100 μg/ml) and protease inhibitor (Roche) and twice with 1x TBS. For IPs taken for western blotting, beads were resuspended in 40 ul 2x SDS buffer and boiled for 5 minutes. For IPs taken for RNA isolation, beads were resuspended in 1x TBS and incubated with 0.5 ul Proteinase K for 15 minutes at 42°C with shaking at 1200 rpm. Supernatant was removed from beads and added to 400 ul Trizol and frozen.

### Western Blotting

For TRAP experiments, input and IP samples were run on an SDS-PAGE gel then transferred to a nitrocellulose membrane. Membranes were blocked in milk for one hour then incubated in 1x TBS containing primary antibody (1:2000 αFLAG (Sigma, M8823) overnight at 4°C. Membranes were washed thrice with 1x TBS then incubated in 1xTBS containing secondary antibody (1:40,000 αMouse HRP) for one hour. Membranes were washed thrice more in 1x TBS before developing with ECL (50% pico/50% femto, Fisher). Blots were imaged on a Bio-Rad ChemiDoc instrument. Band intensities were quantified using ImageJ.

### Bioinformatic Analysis of TRAP-sequencing Data

Reads were aligned to a reference genome (*C. elegans* PRJNA13758 WS275) using STAR (version 2.7.11a). Percent uniquely mapped reads for each sample are listed in **Table S6**. Differential gene expression analyses were performed in R with DESeq2 (Love et al. 2014). Full code for these analyses can be found at: https://github.com/emierdma/GSF3037.

## Data Availability

Strains and plasmids are available upon request. Raw and processed high-throughput sequencing data generated in this study have been submitted to the NCBI Gene Expression Omnibus (GEO; https://www.ncbi.nlm.nih.gov/geo/) under accession number GSE299666 and to the EMBL-EBI European Nucleotide Archive (ENA; https://www.ebi.ac.uk/ena/browser/home) under accession numbers PRJEB100883 and PRJEB100892.

## Acknowledgments

The authors would like to thank Dustin Updike for his guidance with *C. elegans* germline isolation for RNA sequencing, the Indiana University Center for Genomics and Bioinformatics for running the high-throughput sequencing experiments, and the Indiana University Light Microscopy Imaging Center for the use of the Leia SP8 Confocal.

## Supplemental Figure Legends

Figure S1: **Validation of germline-specificity of TRAP method**. IP/input RT-qPCR expression values of germline gene *glh-1* and somatic gene *elt-2* from FLAG IP samples from negative control (wild-type) animals and germline RPL-4::FLAG (HAH87: efls155[*mex-5p*::*rpl-4*::FLAG::*tbb-2* 3’UTR + *Cbr-unc-199*(+)]II) animals. Data represents three biological replicates.

Figure S2: **TRAP pulldowns for TRAP-seq experiment**. Representative western blots (top) of input and IP samples from three biological replicates of TRAP-seq and quantification of IP blots (bottom) relative to wild-type.

Figure S3: **Schematic of germline editing site identification pipeline**.

## Supplemental Tables

Table S1: **Germline editing sites**. Germlines were dissected from young adult wild-type and *adr-2(-)* animals. RNA was isolated from germline samples, poly-A selected, then made into cDNA sequencing libraries and sequenced. Sequencing reads were mapped to a reference genome and SAILOR was used to identify A-to-I editing sites. Editing sites were then narrowed down to sites with ≥99% confidence in all three biological replicates. Editing sites also identified in *adr-2(-)* germlines were removed as false positives. Editing site ID (column A) corresponds to chromosomal coordinate of each site. Column B indicates the genic region of each editing site, Column C indicates Wormbase ID of the gene for each editing site, Column D the corresponding gene name, Columns E-G indicate the percent editing value for each replicate, Column H indicates the average percent editing value. Column I indicates whether each editing site was also identified in whole worms (Schiksnis et al., 2024), Column J indicates whether each editing site was also identified in neurons (Rajendren et al., 2021), Column K indicates whether each edited site occurs in a known maternal transcript (Quarato et al., 2021). For each gene with editing sites only in germlines, Column L indicates whether gene is expressed in young adult germlines and Column M indicates whether the gene is also expressed in young adult neurons (Ghaddar et al., 2023); NA indicates not applicable, ND indicates no data available.

Table S2: **ADR-1-regulated germline editing sites**. Variant calling was performed for all high-confidence germline editing sites (Table S1) on three replicates each of wild-type and *adr-1(-)* germline RNA-sequencing datasets. ADR-1-regulated sites were determined by filtering for sites with >10 reads in all replicates and a >5% change in editing between wild-type and *adr-1(-)* in all replicates. Column A lists the site ID for each edited site, Column B lists the genic region, Column C lists the Wormbase ID of the gene containing each editing site, Column D lists the corresponding gene name. The read coverage for each site is listed in Columns E, G, and I (WT) and K,M, and O (*adr-1(-)*). The % variant reads for each site is listed in Columns F, H, and J (WT) and L,N, and P (*adr-1(-)*).

Table S3: **Differentially expressed genes in *adr-2(-)* and *adr-1(-)* germlines compared to wild-type germlines**. Germlines were dissected from young adult wild-type, *adr-2(-)*, and *adr-1(-)* animals. RNA was isolated from germline samples, poly-A selected, made into cDNA sequencing libraries and sequenced. Sequencing reads were aligned to a reference genome and feature counts followed by DESeq2 was used to identify differentially expressed genes. Differentially expressed genes were narrowed down to those with p-adjusted (padj) ≤ 0.05. Column A indicates the Wormbase ID for each gene, column B indicates the corresponding gene name, Columns C, D, and E indicate the log2 fold change, p-value, and p-adjusted for each gene as determined by DESeq2, Columns F-H indicate expression values from DESeq2 for each of three replicates of *adr-2(-)* germlines, Columns I-K indicate expression values from DESeq2 for each of three replicates of wild-type germlines. Column L indicates the gene ontology category for each misregulated gene.

Table S4: **Differential germline ribosome association of genes in *adr-2(-)* germlines compared to wild-type germlines**. Three biological replicates of TRAP were performed on lysates prepared from young adult HAH87 and HAH88 animals. RNA was isolated from TRAP IP samples, poly-A selected, made into cDNA libraries and sequenced. Sequencing reads were aligned to a reference genome and feature counts followed by DESeq2 was used to identify genes with differential ribosome association (padj ≤ 0.05). Column A indicates the Wormbase ID for each gene, column B indicates the corresponding gene name, Columns C, D, and E indicate the log2 fold change, p-value, and p-adjusted for each gene as determined by DESeq2, Columns F-H indicate counts from FeatureCounts for each of three replicates of *adr-2(-)* TRAP IPs. Columns I-K indicate counts from FeatureCounts for each of three replicates of WT TRAP IPs.

Table S5: **Oligonucleotides used in this study**.

Table S6: **Quality measures for RNA-sequencing**. Related to Figures 1 and 2. Column A lists sequenced samples. Total Sequences (Column B) and Sequences Flagged as Poor Quality (Column C) determined via FASTQC. % uniquely mapped reads (Column D) determined via STAR alignment.

